# Microsecond molecular dynamics of SOD1 variants suggest a structural basis for divergent ALS clinical outcomes

**DOI:** 10.64898/2026.08.29.747999

**Authors:** Abed Alah Refaee, Konstantin Röder, Giancarlo Ruocco, Edoardo Milanetti, Alfredo Iacoangeli

## Abstract

Amyotrophic lateral sclerosis (ALS) is a fatal neurodegenerative disease characterised by progressive motor neuron degeneration. Mutations in the SOD1 gene represent the second most common genetic cause of ALS (ALS), and distinct SOD1 missense variants present with markedly different clinical profiles. A4V leads to an aggressive form of the disease (median survival ∼1y), H46R confers a mild, slowly progressive course and I113T exhibits an intermediate phenotype. The molecular basis by which these mutations produce divergent clinical outcomes remains poorly understood. We performed extensive classical molecular dynamics simulations of wild-type SOD1 and the three ALS-associated variants in the apo monomeric state to attempt to investigate the mechanisms behind such phenotypic differences. Structural stability, global compactness, and conformational flexibility, as well as analysis of collective motions between residues and estimation of free energy, were assessed. The H46R, A4V, and I113T variants exhibited distinct dynamic behaviours, highlighting differences in structural stability, local flexibility, and intramolecular interactions. These findings suggest that specific structural regions may contribute differently to protein dysfunction and could represent key elements for understanding the relationship between molecular dynamic properties and the differing clinical severity associated with these variants. Most strikingly, H46R exhibited exceptional structural stability across every analytical level, the lowest global deviation, most attenuated local flexibility, strongest internal dynamic coordination, and the deepest, most confined free energy basins of any system examined. This convergent multi-layered evidence of structural restraint provides a compelling mechanistic basis for the mild and slowly progressive clinical course of H46R ALS, suggesting that enhanced conformational rigidity, rather than bulk destabilisation, is the defining biophysical feature of this variant, and that its pathogenic mechanism operates through a route fundamentally decoupled from the aggregation-driven toxicity that characterises the more aggressive SOD1-ALS mutations. mutations.

## 1. Introduction

Amyotrophic lateral sclerosis (ALS) is a progressive and invariably fatal neurodegenerative disease characterised by the selective degeneration of upper and lower motor neurons.^1,2^ Affected individuals typically present with focal weakness that spreads progressively, leading to paralysis and death from respiratory failure within 2– 5 years of symptom onset. While 90% of ALS cases are sporadic, approximately 10% are familial (fALS).^3^ Mutations in the SOD1 gene were first identified in 1993 and account for approximately 10–15% of fALS and 1–2% of all ALS cases.^4,5^

SOD1 is a 32 kDa homodimeric metalloenzyme that catalyses the dismutation of superoxide radicals^6,7^. The physiologically active form is a fully metalated homodimer (holo-SOD1) in which each 153-residue monomer adopts a Greek key *β*-barrel fold^8–9^ with structural integrity depending on copper and zinc coordination within the metal-binding loop (MB Loop; loop IV, residues 49–83), a Cys57–Cys146 disulfide bond, and homodimerisation.^10–12^ The electrostatic loop, (E Loop, loop VII, residues 121–142) gates substrate access to the catalytic copper site (Figure 1A).^10^ In the context of ALS pathogenesis, however, metal loss and disulfide reduction produce the apo monomeric form, the immature, partially unfolded precursor state that is directly implicated in misfolding and aggregation.¹⁰^˒^¹²^˒^¹³ It is this pathologically relevant apo monomeric state that is modelled in the present study, as it represents the conformational species most prone to the aberrant self-association underlying motor neuron toxicity in SOD1-ALS.

**Figure 1.**
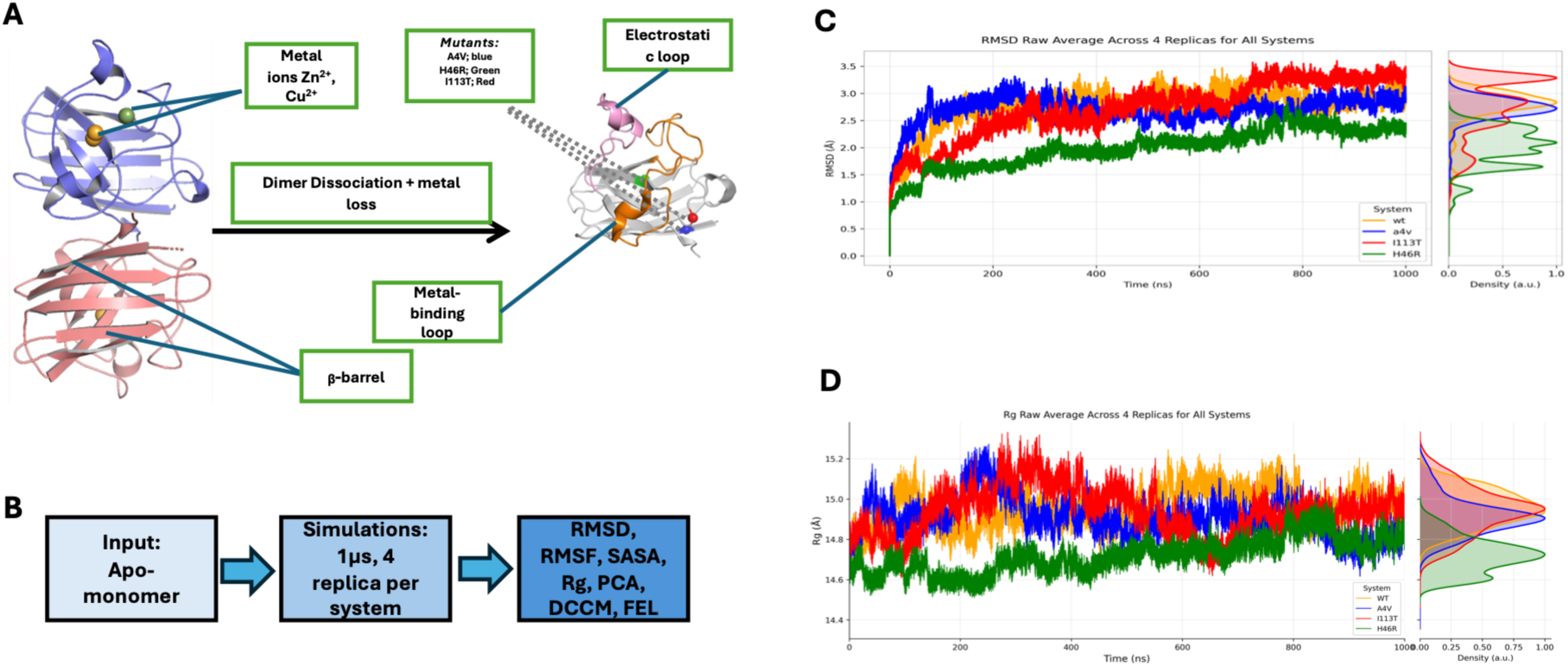
Structure and global dynamics of wild-type SOD1 and ALS-associated mutants over 1 μs molecular dynamics simulations. **(A)** Left: representation of the SOD1 homodimer (PDB: 2C9V) with Chain 1 (slate blue) and Chain 2 (magenta) shown in cartoon. Copper (Cu²⁺, orange spheres) and zinc (Zn²⁺, green spheres) ions are shown in their respective binding sites. The Greek key β-barrel fold, metal-binding loop (residues 49–83), and electrostatic loop (residues 121–142) are annotated. Right: the apo-monomer form following dimer dissociation and metal loss, with the metal-binding loop and electrostatic loop, and point mutations highlighted. **(B)** Simulation workflow: apo-monomer structures were subjected to 1 μs MD simulations with four independent replicas per system, followed by analysis of RMSD, RMSF, SASA, Rg, PCA, DCCM, and FEL. **(C)** Backbone RMSD averaged across n = 4 independent replicas. H46R plateaus at the lowest deviation (∼2.2–2.5 Å); I113T reaches the highest (∼3.0–3.5 Å); A4V and WT occupy an intermediate, overlapping range. **(D)** Radius of gyration (Rg). H46R is consistently the most compact (∼14.6–14.8 Å) with a bimodal KDE distribution indicative of two-state sampling; WT, A4V, and I113T occupy a higher, overlapping range (∼14.8–15.2 Å), with I113T showing the broadest spread. Right-hand panels in (C) and (D) show marginal Gaussian kernel density estimates. Colour scheme: WT (orange), A4V (blue), H46R (green), I113T (red).

More than 200 distinct SOD1 mutations produce markedly different disease phenotypes.^13,14^ Work from our group and others has established a decoupling between mechanisms governing disease risk and those determining duration ^15^. Three variants exemplify this heterogeneity: A4V causes aggressive disease (median survival approximately 1 year) through cytotoxic aggregation and dimer-interface destabilisation;^9^ H46R abolishes copper binding yet confers a slow disease course of a decade or more and has a relatively low propensity to form classical insoluble amyloid aggregates, and I113T, a beta-barrel packing defect, exhibits variable penetrance. ^14,44^

Experimental studies have characterised thermodynamic stability and aggregation propensity of SOD1 variants but offer limited insight into residue-level dynamics in real time.^16,17^

We previously investigated these variants by molecular dynamics simulations, providing early evidence that distinct structural mechanisms underlie divergent clinical phenotypes;^15^ however, the simulation timescales were insufficient to adequately sample the conformational landscape of the apo monomeric state. Here, we address this limitation by extending the sampling of the system to achieve a more comprehensive characterisation of its structural dynamics.

In this work, we use a hierarchical and progressively detailed analysis framework. We first characterize the global structural properties of the apo monomeric protein, the pathologically relevant precursor to misfolded and aggregated species.^11,18,19^ In our analysis, we include root mean square deviation (RMSD) and radius of gyration (Rg), to assess overall stability and compactness. We then move to a residue-level analysis, examining solvent-accessible surface area (SASA) and residue-wise fluctuations over the simulation time (RMSF), in order to capture local structural rearrangements induced by each mutation. Furthermore, by analysing residue–residue correlated motions through dynamic cross-correlation (DCCM) analysis and principal component analysis (PCA), we investigate collective and synergistic motions across the protein and characterize how each pathogenic mutation affects sampling of the essential conformational space defined by the dominant principal components. Additionally, the study aims to quantify the thermodynamic basis of the observed conformational behaviour through free-energy landscape (FEL) reconstruction, providing a comprehensive energetic description of mutation-induced structural effects.

## 2. Materials & Methods

### 2.1 System preparation and starting structures

Four simulation systems were prepared: WT (PDB: 2C9V) ^20^, A4V (PDB: 1UXM)^21^, H46R (PDB: 1OZT), ^22^ and I113T (PDB: 1UXL).^23^ Using Pymol 3.1.6.1, ^24^ for each system, a single monomer was extracted; all metal ions (Cu^2+^, Zn^2+^) and crystallographic waters were removed, as we aim to study the dynamics of the apo-monomer, a species implicated in misfolding and aggregation. PDBFixer ^45^ was used to check for missing atoms, bonds, alternative locations and chirality. Hydrogen addition was performed using gmx pdb2gmx.

### 2.2 Molecular dynamics simulation protocol

All simulations used GROMACS 2021.5 ^25,26^ with the CHARMM27 ^27^ all-atom force field (ff), a well-validated all-atom protein force field with a long track record in SOD1 and other beta-barrel protein simulations.^28,29^ Each system was placed in a dodecahedral box (minimum 1.0 nm protein-to-edge), solvated with SPC216 water, the recommended model for CHARMM ff and neutralised with Na+ or Cl− counter-ions, to achieve net zero charge, as required for the particle mesh Ewald (PME) treatment of long-range electrostatics under periodic boundary conditions. ^30,31^ Energy minimisation used the steepest descent algorithm until forces converged below 1000 kJ mol⁻¹ nm⁻¹. Equilibration comprised 100 ps NVT at 300 K using Berendsen thermostat followed by 100 ps NPT at 1 bar Parrinello–Rahman barostat with position restraints on all heavy atoms. Production runs of 1 μs per replica used a 2 fs timestep, LINCS bond constraints, 1.0 nm non-bonded cutoff, and PME electrostatics.^31–34^ Four independent replicas per system yielded 16 μs total aggregate sampling.

### 2.3 Global structural stability analysis

Backbone RMSD, total SASA, and Rg were computed over the full 1 μs production trajectories using the GROMACS tools with default settings. Time series were averaged across n = 4 independent replicas. Marginal probability distributions were estimated for each observable by fitting a Gaussian distribution to the replicate-averaged time series, with the probability density function:

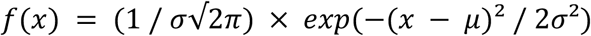

where μ and σ are the sample mean and standard deviation of the observable respectively. Distributions were visualised as histogram-overlaid Gaussian fits using matplotlib and are shown in the right-hand panels of Figure 1C-D. ^35^

### 2.4 Local flexibility analysis

RMSF and SASA were obtained using the GROMACS tools with default settings, averaged across all four replicas. Loop-specific RMSD was computed independently for the E-loop (residues 121–142) and MB-Loop (residues 49–82) relative to their equilibrated conformations with restricted index groups.

### 2.5 Principal component analysis & DCCM

PCA was performed on C-alpha coordinates from equilibrated trajectory portions. The covariance matrix was diagonalised using GROMACS PCA utility. PC1, PC2, and PC3 were retained. Each trajectory was projected onto the shared eigenvector basis derived from the concatenated four-system trajectory.

DCCMs were computed from concatenated, fitted, multi-replica trajectories (4×1 μs; subsampled every 10th frame, approximately 40,000 frames per system). The cross-correlation coefficient Cij between residues i and j was calculated as:

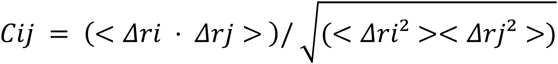

where *Δri* and *Δrj* are C-alpha displacement vectors from time-averaged positions. *Cij* ranges from +1 (fully correlated) through 0 (uncorrelated) to −1 (fully anti-correlated). Rigid-body motions were removed by least-squares fitting prior to computation. DCCMs were computed using Bio3D ^36^ and visualised as contour-filled heatmaps.

### 2.6 Free energy landscape reconstruction

Two-dimensional FELs were reconstructed from PCA projections onto the shared eigenvector basis (gmx anaeig). The Gibbs free energy surface was computed as:

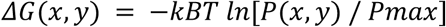

where *P*(*x*, *y*) is the probability density from a two-dimensional histogram (100 bins per dimension), *Pmax* is the maximum bin density, *kB* is the Boltzmann constant, and T = 300 K. Unpopulated bins were capped at 14 kJ/mol. Two projections were retained: PC1 vs PC2 and PC1 vs PC3.

### 2.7 Visualisation and analysis tools

Structural visualisations were generated using Pymol 3.1.6.1. ^24^ Data plots and analyses were produced using Python with the pandas, NumPy, and matplotlib libraries.^35, 37,38^

## 3. Results

### 3.1 Global structural stability: RMSD and Rg

Backbone root mean square deviation (RMSD) measures the average displacement of backbone atoms from a reference structure over time, providing a global measure of structural stability and the extent to which the protein deviates from its starting conformation. Whereas, the radius of gyration (Rg) describes the mass-weighted distribution of atoms about the protein’s centre of mass, providing a measure of overall structural compactness and the degree to which the protein adopts an extended or globular conformation. Together these metrics quantify structural deviation from the starting conformation, and overall compactness respectively, providing a two-layer portrait of global dynamic behaviour independent of local residue-level features.

#### 3.1.1 Wild-type SOD1 apo monomer as a destabilised reference

WT displayed RMSD in the range 2.8–3.2 Å, and Rg comparable to A4V and I113T (Figure 1C-D). This is expected: the holo-dimeric crystal structure represents the mature, fully metalated form of the protein, whereas the apo monomer simulated here lacks both metal cofactors and the stabilising contacts of the dimer interface. The WT apo monomer therefore serves as a destabilised reference state against which mutation-specific perturbations are assessed (Figure 1A). Its global metrics are comparable to those of A4V and I113T, indicating that monomerisation and metal removal alone impose substantial structural deviation before any mutation-specific effects are considered.

#### 3.1.2 H46R is the most structurally stable and compact variant

H46R was the most stable and restrained system across all three global metrics. RMSD plateaued at a range of 2.2–2.5 Å (Figure 1C), substantially below all other systems. Rg was the most compact of any system (14.6–14.8 Å). Notably, the Rg distribution was bimodal, with two well-separated peaks at approximately 14.6 Å and 14.7Å, suggesting that H46R preferentially samples two distinct global conformational substates. This two-state behaviour is interrogated in 3.2 to further understand residue level drivers of this differential Rg pattern.

#### 3.1.3 I113T exhibits the greatest conformational heterogeneity

I113T displayed the highest global deviation across all metrics: RMSD reached 3.0–3.5Å with a broad right-skewed distribution (Figure 1C). The high-SASA tail is consistent with transient exposure of normally buried surface in a subset of sampled conformations; per-residue SASA decomposition to identify the specific residues contributing to this tail is presented in Section 3.2 (Figure S[1]). The Ile113Thr substitution is located at a buried beta-barrel position; whether this substitution significantly disrupts the local hydrophobic packing contacts at position 113 is examined directly from the simulation data in Section 3.2, where residue-level analyses are presented (Figure 2).

**Figure 2.**
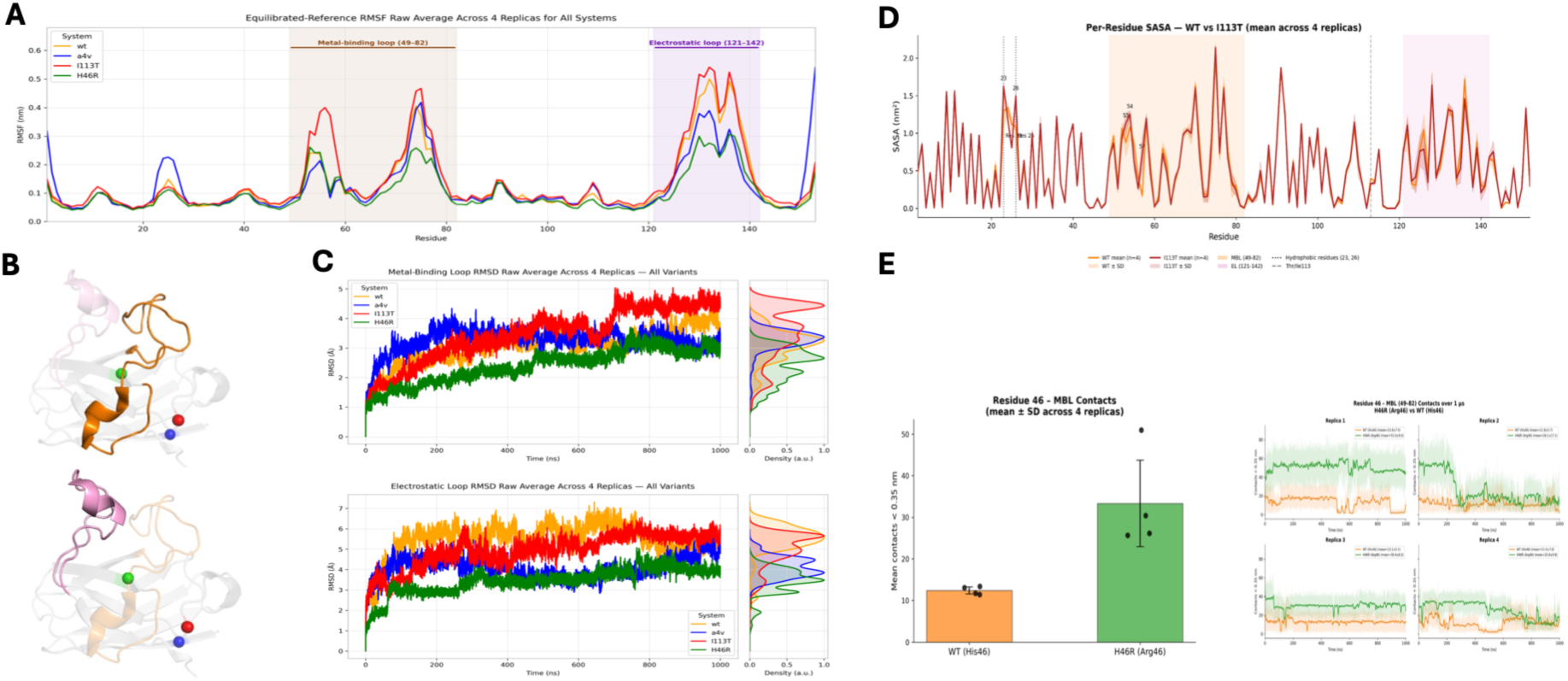
*Local loop dynamics of wild-type SOD1 and ALS-associated mutants over 1 μs MD simulation. **(A)** Per-residue RMSF with regional annotations. Metrics averaged across n = 4 replicas. **(B)** Top. Structural representation of MB Loop. Bottom. Structural representation of E Loop RMSD **(C)** Top. MB Loop RMSD. Bottom. E Loop RMSD **(D**)* Per-residue solvent-accessible surface area (SASA) comparison between wild-type SOD1 and I113T averaged across n = 4 independent replicas. Mean SASA (nm²) per residue for WT (orange) and I113T (red); shaded bands indicate ± SD across replicas. Residues 1 and 153 are excluded. MB Loop and E Loop are highlighted in orange and pink respectively; the mutation site (Ile/Thr113) is indicated by a grey dashed line; hydrophobic residues of interest (Val23, Ile26) are marked by black dotted lines. I113T shows the greatest elevation at Val23 (+0.338 nm², +26.2% relative to WT) and Ile26 (+0.418 nm², +38.6%), consistent with increased solvent exposure of normally buried hydrophobic β-barrel surface. Notably, several E Loop residues (136, 126, 135) and MBL residues (55, 56, 63) exhibit modestly lower SASA in I113T than WT, indicating that the mutation does not uniformly increase solvent exposure across the protein but rather induces a redistribution of surface accessibility, with hydrophobic β-barrel residues becoming more exposed while certain loop surface residues become relatively more buried. ***(E)*** Right. Number of heavy atom contacts within 0.35 nm between residue 46 and MB Loop backbone atoms per replica; solid lines show 5 ns running averages, shaded regions show raw values. **(E)** Left, Mean ± SD contact count across n = 4 independent replicas with individual replica values overlaid. Arg46 (H46R, green) maintains 33.3 ± 10.4 contacts compared to 12.4 ± 0.8 for His46 (WT, orange), consistent with compensatory MBL stabilisation by the guanidinium group

#### 3.1.4 A4V shows rapid destabilisation at an intermediate level

A4V equilibrated to an intermediate global RMSD (2.8–3.0 Å), comparable to WT over much of the trajectory, but reached this plateau within the first 100–200 ns, substantially faster than WT (Figure 1C). Rg overlapped broadly with WT and I113T, confirming that bulk compactness is not dramatically altered at the global level (Figure 1D). The convergence of A4V global metrics with those of WT, despite its dramatically worse clinical prognosis (median survival approximately 1 year), underscores the fundamental limitation of global stability measures in stratifying clinical severity and motivates the finer-grained local and collective analyses presented in subsequent sections.

### 3.2 Local loop dynamics: electrostatic loop and metal-binding loop

The E Loop and MB Loop govern substrate channelling and metal cofactor coordination and are liberated from stabilising constraints in the apo monomeric state, making their dynamics particularly sensitive to mutation.^10^ Per-residue root mean square fluctuation (RMSF) measures the time-averaged positional variability of each residue’s Cα atom over the trajectory, providing a local measure of flexibility that identifies which regions of the protein are most dynamic. Solvent-accessible surface area (SASA) quantifies the total protein surface area accessible to solvent molecules, serving as a measure of surface exposure and the degree to which hydrophobic core residues are exposed. SASA, Loop-specific RMSD and RMSF as well as contact analysis profiles are shown in Figure 2.

#### 3.2.1 The E loop is most deviated in WT and I113T

The E Loop displayed the largest absolute RMSD values of any region examined. WT reached the greatest deviation (6.0-7.0 Å; Figure 2C), underscoring that apo monomerisation alone substantially destabilises this loop independent of mutation. I113T tracked closely behind (5.5-6.2 Å), consistent with the elevated E Loop RMSF peak at residue approximately 57 (Figure 2C). A4V occupied an intermediate range (3.5-5.2 Å) with a bimodal distribution. H46R was markedly attenuated (3.0-4.2 Å) also with a bimodal distribution, suggesting two-state EL sampling consistent with the two Rg substates identified in Section 3.1.2 (Figure S [5]).

#### 3.2.2 I113T drives the greatest MB loop deviation

I113T emerged as the most deviated system at the MB Loop (RMSF 4.3–5.0 Å; broadest distribution; Figure 2A). This is mechanistically notable: Ile113 is located within the β-barrel core, distal from the MB Loop, yet its effects propagate to destabilise the zinc-coordinating loop, indicating long-range allosteric coupling between the β-barrel scaffold and the MB Loop.

The I113T SASA was the most elevated and variable of any system (88×10^2^–92×10^2^ Å²) with a pronounced high-SASA tail (Figure S[1]). Per-residue SASA analysis (Figure 2D) corroborates this: MBL residues 70 (Leu; 1.61×10^2^ Å²), 75 (Gly; 2.14×10^2^ Å²), and 77 (Gly; 1.60×10^2^ Å²) show the highest loop exposure, with the hydrophobic Leu70 being of particular interest given its role in MB Loop packing. Elevated E Loop exposure is also evident at residues 128 (1.64×10^2^ Å²) and 136 (1.46 x10^2^ Å²), consistent with the amplified MB Loop–E Loop anti-correlated dynamics identified by DCCM (Section 3.3). Alongside several β-barrel residues (residues 9, 11, 23, and 26; 1.50–1.63 x10^2^ Å²). Notably, residues 23 (Val), and 26 (Ile) are β-strand hydrophobic and normally buried residues, suggesting that their differentially elevated SASA in I113T reflects transient exposure of normally sequestered hydrophobic surface, a finding consistent with the elevated aggregation propensity of this variant (Figure 2D, Supplementary Figure S [2]).

#### 3.2.3 A4V is distinguished by terminal flexibility at the dimer interface

RMSF profiles revealed a signature unique to A4V: pronounced spikes at the N-terminus (residues 1–5, approximately 3.2 Å) and C-terminus (residue approximately 153, approximately 5.4 Å), both substantially elevated above all other systems (Figure 2A). These regions form the primary dimer interface, implicating enhanced self-association propensity as a driver of A4V toxicity. At the EL and MBL, A4V RMSF largely co-varies with WT, confirming that the dominant local perturbation in A4V is terminal rather than loop-centred, mechanistically distinct from I113T, whose primary local signature is at the EL and MB Loop.

#### 3.2.4 H46R displays uniformly attenuated local flexibility underpinned by compensatory Arg46–MBL interactions

H46R consistently exhibited the lowest fluctuations across all local metrics, with the RMSF trace running below all other systems at nearly every residue position (Figure 2A). Despite carrying the mutation at His46, a direct zinc ligand, H46R displayed the most restrained MB Loop dynamics (2.8–3.3 Å) of any system (Figure 2C). H46R’s SASA remained narrow and low (8500–8700 Å²) with a unimodal distribution (Figure S[1]).

Contact analysis between residue 46 and the MB Loop reveals a structural basis for this paradox: Arg46 maintained 33.3 ± 10.4 contacts with MB Loop atoms over the 1 μs trajectory, compared to 12.4 ± 0.8 for His46 in WT (Figure 2E). The guanidinium group of Arg46 forms more numerous, shorter, and more geometrically optimal hydrogen bonds with MBL backbone atoms than His46 (Supplementary Figure S[4]), effectively compensating for the loss of zinc coordination by mechanically constraining the loop. The uniform suppression of E Loop, MB Loop, and terminal flexibility reinforces the global restraint observed in Section 3.1 and supports a pathogenic mechanism for H46R that is decoupled from the dynamic destabilisation characteristic of A4V and I113T. That H46R nonetheless causes ALS suggests mechanisms beyond bulk destabilisation, such as altered metal ion affinity or impaired dimer reassembly.

Structural comparison of representative frames extracted from each Rg peak, shown in Figure1, revealed that the two substates differ predominantly in the position of the E Loop (residues 121–142), with the metal-binding loop and beta-barrel core remaining largely invariant between them (Supplementary Figure S [5]). This two-state behaviour is consistent with the bimodal E Loop RMSD distribution observed in Figure 2C.

### 3.3 Principal component analysis of collective motions

Principal component analysis (PCA) of Cα atomic coordinates identifies the dominant collective motions of the protein by diagonalising the covariance matrix of positional fluctuations, yielding a set of orthogonal eigenvectors (principal components) that describe the directions of greatest variance in conformational space; projecting each trajectory onto these components allows the conformational ensemble sampled by each variant to be visualised and compared in a reduced-dimensional space.

Systems’ trajectories were concocted, rotations and translations removed, and C-ɑ trajectory extracted. PCA was performed on C-ɑ coordinates using a shared covariance matrix from the concatenated four-system trajectory. PC1-PC3 collectively captured approximately 49% of conformational variance, with PC1 and 2 most dominant explaining 20.4% and 19.6% of the variance respectively, while PC3 only 8.3% of the variance (Figure 3A). Per-residue loadings (Figure 3B-D) identified the structural drivers of each component: PC1 is dominated by the E loop peak contribution approximately 0.4; PC2 reflects coupled rocking between the zinc-binding loop (residue 63-82) and E loop; PC3 is governed by the zinc-binding loop and C-terminal region. Putty representations of each component are shown inside Figures 3B-D.

**Figure 3.**
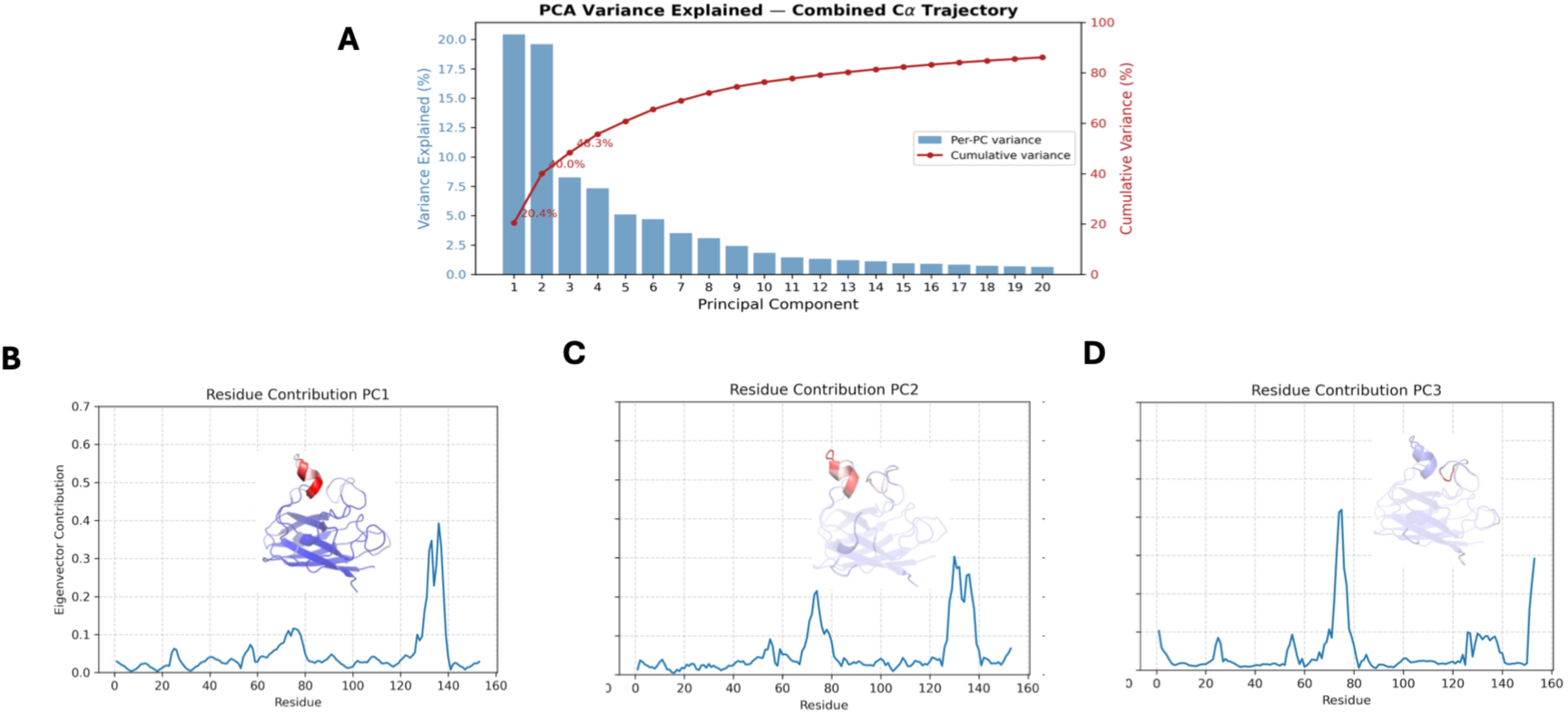
Principal component analysis of collective motions in wild-type and mutant SOD1. **(A)** Variance explained by the first ten principal components of the shared C-alpha eigenvector basis. **(B–C)** Per-residue loadings for PC1–PC3. Electrostatic loop dominates PC1; zinc-binding and electrostatic loops co-dominate PC2; zinc-binding loop and C-terminal region dominate PC3. Structures with residues coloured according to the residue contributions are overlayed above curves with colour (Blue-Red) indicating the degree of region contribution to PC.

#### H46R

H46R shifted substantially along PC1 toward negative values relative to WT, reflecting altered electrostatic loop dynamics, while PC2 sampling remained broadly comparable (Figure S[3], row 1).

#### A4V

Despite extensive overlap with WT in the PC1–PC2 plane, A4V accessed a well-defined conformational basin in the PC1–PC3 subspace (PC1, PC3 approximately (−1,−1) to (+2,−2.5)) not substantially populated by WT (Figure S[3], row 2).

#### I113T

I113T showed the greatest conformational similarity to WT, with near-superimposable PC1–PC2 densities. A modest separation emerged only in the PC2–PC3 plane (Figure S[3], row 3), consistent with its slower progression and variable penetrance.

PCA revealed a gradient of conformational perturbation: H46R shifted along PC1, mainly characterised by E Loop; A4V accessed a distinct PC1-PC3 basin; I113T showed subtle PC2-PC3 redistribution. Full pairwise cross-projections are provided in Supplementary material (Figure S[3]).

##### 3.3.1 DCCM

DCCM adds a directional dimension inaccessible from magnitude-based metrics, mapping whether residue pairs move together (Cij > 0) or in opposition (Cij < 0) (Figure 4A-D; Table 4E).

**Figure 4.**
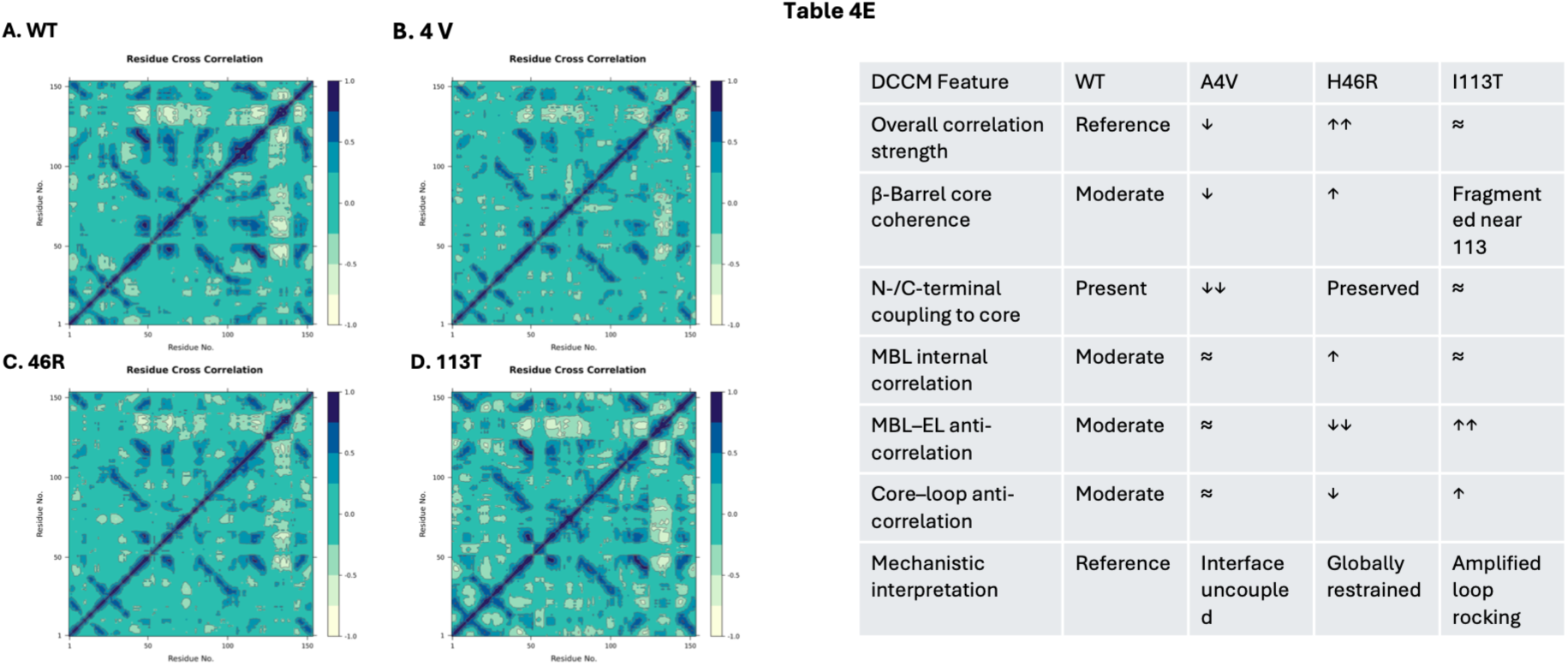
Dynamic cross-correlation matrices (DCCMs) of wild-type SOD1 and ALS-associated mutants. Pairwise C-alpha cross-correlation coefficients (Cij) computed from concatenated multi-replica trajectories (4 x 1 μs, subsampled every 10th frame). Positive values (dark blue) indicate correlated motions; negative values (yellow/light) indicate anti-correlated motions. (**A**) WT, (**B**) A4V, (**C**) H46R, (**D**) I113T. **Table 4.D** DCCM analysis of WT and SOD1 mutants. Arrows indicate changes in correlation relative to WT: ↑ (increase), ↓ (decrease), ≈ (similar).

The MB Loop–E Loop axis (residues 49–83 vs 121–142) was the most variant-sensitive region. WT displayed moderate anti-correlated rocking between these loops, consistent with the coupled PC2 loading. H46R markedly suppressed this anti-correlation, locking both loops and reducing the large-amplitude opposing excursions that could expose aggregation-prone surfaces. I113T showed the strongest MB Loop– E Loop anti-correlation of all systems despite the mutation site being distal from both loops, confirming allosteric amplification through the beta-barrel core (residues 85– 115). A4V preserved MB Loop–E Loop anti-correlations at near-WT levels.

At the dimer interface (residues approximately 1–10 and approximately 140–153), A4V was sharply distinguished: while WT, H46R, and I113T maintained correlated terminal–core coupling, A4V showed fragmented and weakened correlations. The A4V termini do not merely fluctuate more, they move independently of the scaffold, consistent with interface residues primed for aberrant self-association.

H46R was globally the most internally coordinated system, with the broadest near-diagonal correlated band and enhanced MBL coherence despite loss of zinc coordination, providing a mechanistic rationale for its resistance to the aggregation pathway.

In summary, DCCM reveals three qualitatively distinct mechanisms of dynamic reprogramming: interface uncoupling (A4V), global dynamic restraint (H46R), and amplified loop–loop anti-correlations (I113T), distinctions not resolvable from fluctuation amplitudes alone, and a natural basis for the thermodynamic stratification presented in the FEL analysis below.

### 3.4 Free energy landscapes

The FEL integrates the preceding structural and directional analyses into a single thermodynamic picture (Figure 5). In the PC1–PC2 projection, H46R occupied the most confined landscape, with three discrete deep minima clustered within a narrow PC space (PC1 approximately 0 to +2) and steep surrounding barriers, consistent with its globally restrained dynamics. WT displayed the broadest and most diffuse surface, spanning PC1 approximately −4 to +2 with multiple shallow, loosely connected minima. A4V showed a more compact landscape shifted toward positive PC1/PC2. I113T rivalled WT in breadth, extending further along negative PC2 (approximately −3) with multiple minima of comparable depth.

**Figure 5.**
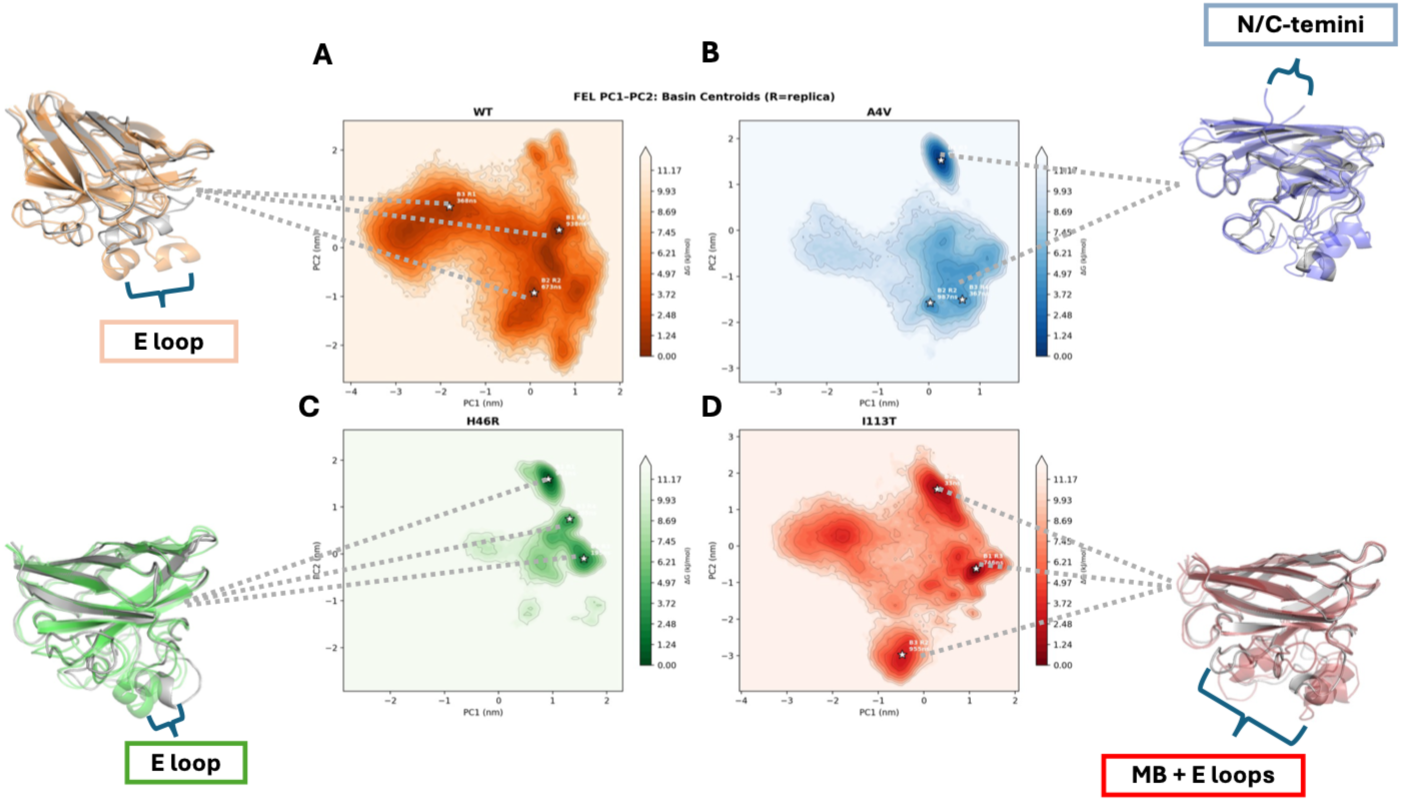
Free energy landscapes and basin conformational ensembles of wild-type SOD1 and ALS-associated mutants. **(A-D)** Two-dimensional free energy surfaces (ΔG = −kBT ln P/Pmax) in the PC1–PC2 plane reconstructed from projections onto a shared eigenvector basis. Stars indicate basin centroids: PC1–PC2 free energy landscapes. Darker colours indicate lower free energy; white indicates unpopulated regions (capped at 14 kJ/mol). Stars indicate identified basin centroids. Superimposed basin centroid structures for **B. WT** basins (B1– B3). Grey: global minimum (B1, R4, 938 ns); transparent orange overlays: B2 (R2,674 ns) and B3 (R1,368 ns). **C**. **A4V** basins minimum (B1, R3, 925 ns); transparent blue overlays: B2 (R2, 987 ns) and B3 (R4,367 ns). **D. H46R** basins (B1–B3). Grey: global minimum (B1, R1, 921 ns); transparent green overlays: B2 (R3,195 ns) and B3 (R4,205 ns). **E. *113T*** *basins (B1–B3). Grey: global minimum (B1, R3, 746 ns); transparent red overlays: B2 (R4,33 ns) and B3 (R2,955 ns)*

Basin centroid structures (Figure 5 A–D) reveal the structural character of inter-basin displacement within each system. WT inter-basin variation is diffused across the MB Loop and E Loop without a dominant shift. In A4V, secondary basins show marked N- and C-terminal displacement relative to the global minimum, reinforcing the DCCM finding of terminal uncoupling. H46R basin centroids are structurally similar, with inter-basin differences localised to subtle E Loop reorientations. I113T centroids span the broadest conformational range, with inter-basin displacement distributed across both the MB Loop and E Loop, consistent with the amplified MB Loop–E Loop anti-correlations and low interconversion barriers.

The PC1–PC3 projection (Supplementary Figure S[7]) provided additional discrimination: H46R collapsed into a single narrow basin; A4V populated a well-defined minimum at PC1 approximately 0, PC3 approximately −0.5 to −1.5, representing a thermodynamically stable intermediate not accessible to WT; and I113T exhibited at least three shallow minima with low interconversion barriers.

Taken together, the FEL provides a thermodynamic rationale for each variant’s clinical behaviour: the deep narrow basin of H46R explains its heightened structural stability with no thermodynamically stable aggregation-prone intermediates; the distinct A4V intermediate may serve as a kinetic gateway to toxic aggregation; and the broad multi-minima landscape of I113T provides a structural basis for variable penetrance.

## 4. Discussion

Microsecond-scale MD simulations of the SOD1 apo monomer resolved variant-specific dynamic signatures for A4V, H46R, and I113T that suggest a coherent map onto their divergent clinical phenotypes. The layered analytical framework, spanning global stability, local flexibility, collective motions, directional coupling, and thermodynamic landscape reconstruction, demonstrates that each variant perturbs the apo monomer through a mechanistically distinct route, with each analytical layer revealing features inaccessible to the preceding one.

### 4.1 Relation to prior computational work on SOD1-ALS

Although this study builds on previous molecular dynamics work of our group on ALS SOD1 variants ^15^ that proposed a decoupling between the structural mechanisms governing the decoupling between ALS onset and progression observed in our large multi-centre retrospective observational study. The present work identifis loop-region dynamics as a key discriminating feature offering a new design that corroborates these findings at substantially extended timescales (1 μs per replicate, 4 replica per system) and adds DCCM and FEL as analytical layers, resolving the distinction between increased fluctuation amplitude and loss of directional coordination, a dimension inaccessible to per-residue flexibility measures alone. The mechanistic classification proposed here provides thermodynamic and network-level grounding for the phenotypic heterogeneity previously identified.

### 4.2 Cross-experiment variability and simulation bias

Each variant was initialised from a distinct experimental structure, introducing potential biases from differences in crystallographic resolution, or crystal contacts. All systems underwent identical equilibration as apo monomers with metal ions removed. Critically, global stability metrics were replicated in parallel simulations where all three variants were introduced via in-silico point mutation into the WT reference structure (PDB: 2C9V), confirming robustness to starting-structure choice (Figure S[6]). Experimental structures were retained for the full analysis to preserve native side-chain geometries; the concordance between both sets constitutes internal validation of the reported dynamic signatures.

### 4.3 H46R: structural restraint and a decoupled pathogenic mechanism

H46R was the most structurally stable and internally coordinated system at every analytical level: lowest RMSD, narrowest SASA, most compact Rg, most attenuated loop flexibility, and the deepest narrow FEL basin. Despite abolishing zinc coordination, the Arg46 guanidinium group appears to compensate by locking the MB Loop into a coherent dynamical unit and suppressing the MB Loop–E Loop anti-correlated rocking characteristic of WT, thereby reducing the probability of populating misfolding-competent conformations. ^39^ The slow clinical progression of H46R is therefore consistent with a pathogenic mechanism decoupled from bulk destabilisation,^15^ with impaired copper delivery or aberrant chaperone interactions more likely to predominate.

### 4.4 A4V: interface uncoupling and a thermodynamically distinct aggregation-prone intermediate

Global metrics placed A4V close to WT, reinforcing the observation from our previous work that bulk stability measures alone are insufficient to stratify clinical severity ^15^. The pathogenic signature emerged at finer scales: terminal RMSF spikes at the dimer interface, a novel conformational basin in PC1–PC3 space, and DCCM-resolved uncoupling of the termini from the beta-barrel core. The FEL confirmed this basin as a thermodynamically stable intermediate rather than a transient excursion, an energetically accessible misfolded state that may serve as a kinetic gateway to the toxic aggregation characteristic of A4V ^40,41^.

### 4.5 I113T: amplified loop anti-correlations and variable penetrance

Despite a mutation site distal from both functional loops, I113T exhibited the largest MB Loop deviation, elevated E Loop flexibility, and the strongest MB Loop–E Loop anti-correlation of all four systems, a gain-of-function in opposing loop dynamics propagated through the beta-barrel core. ^42,43^ The FEL revealed at least three low-barrier minima, indicating superficial interconversion between native-like and destabilised states. Whether ALS develops in I113T carriers, and at what rate, may depend on the population balance of this conformational equilibrium, which is plausibly sensitive to genetic background and cellular stress, a structurally grounded basis for the variable penetrance that defines this variant clinically.

### 4.6 A mechanistic framework for SOD1-ALS phenotypic heterogeneity

The results support a model in which three ALS mutations perturb the SOD1 apo monomer through distinct biophysical routes. A4V destabilises the dimer interface through terminal uncoupling, populating a thermodynamically stable aggregation-prone intermediate consistent with its aggressive clinical course. H46R, paradoxically, enhances internal dynamic coordination and confines the protein to a deep narrow energy basin, producing slow non-aggregation-dependent disease. I113T propagates a beta-barrel packing defect to amplify MB Loop–E Loop anti-correlated dynamics and broaden the conformational landscape, generating the thermodynamic plasticity that underlies variable penetrance. This stratification is consonant with the onset– progression decoupling proposed previously by our group^15^ and may set the basis of a framework for interpreting the broader landscape of SOD1 mutations.

### 4.7 Limitations and future directions

Several limitations merit acknowledgement. The fixed-charge CHARMM27 force field cannot capture electronic polarisation or explicit metal coordination chemistry. At 1 μs per replica, rare transitions between FEL basins may not be fully converged; enhanced sampling methods such as replica exchange or meta-dynamics would strengthen these findings. Only the monomeric apo state was simulated, excluding dimer dissociation kinetics, molecular crowding, and chaperone interactions. FEL reconstruction from PCA projections is limited to linear collective motions. Future work should extend the mechanistic classification to additional clinically characterised SOD1 variants, incorporate network community analyses to resolve allosteric pathways, and validate computational predictions against experimental aggregation kinetics.

## 5. Conclusions

Using standard descriptors, including RMSD, SASA, and Rg, complemented by analyses of residue-level fluctuations and collective motions. In addition, dynamic cross correlation matrix (DCCM) and conformational free-energy landscapes (FEL) were evaluated to characterize the structural impact of pathogenic mutations. H46R was the most structurally stable variant across structural stability and global compactness measures with uniformly attenuated local flexibility. A4V showed pronounced N- and C-terminal flexibility implicating dimer-interface destabilisation. I113T displayed the greatest metal binding loop (MB-Loop) deviation and elevated Electrostatic Loop (E-Loop) flexibility consistent with allosteric propagation from the disrupted **β**-barrel core. PCA, DCCM, and FEL analyses revealed mutation-specific dynamic signatures at every analytical level. Variant-specific dynamic fingerprints rationalise the divergent clinical outcomes of A4V, H46R, and I113T at the atomic level, identifying the dimer interface, electrostatic loop, and metal-binding loop as key structural determinants of phenotypic divergence.

Together, these results demonstrate that a layered MD framework, integrating global stability, local flexibility, PCA, DCCM, and FEL, can resolve variant-specific dynamic fingerprints of the SOD1 apo monomer and provide a structurally grounded basis for interpreting genotype–phenotype relationships in SOD1-ALS. The dimer interface, electrostatic loop, and metal-binding loop emerge as the principal structural determinants of phenotypic divergence and as candidate targets for future therapeutic and mechanistic investigation.

## Acknowledgements & Funding

The first author’s work was supported by the UK Engineering and Physical Sciences Research Council (EPSRC) [Grant reference number EP/Y035216/1] Centre for Doctoral Training in Data-Driven Health (DRIVE-Health) at King’s College London, with additional support from the National Institute for Health and Care Research (NIHR) Maudsley Biomedical Research Centre (BRC) [Grant reference number NIHR203318]. The views expressed are those of the author and not necessarily those of the NHS, the NIHR or the Department of Health and Social Care. Simulations and analyses were conducted using King’s College London high-performance computer King’s Computational Research, Engineering and Technology Environment (CREATE). https://doi.org/10.18742/rnvf-m076.

Corresponding author and first supervisor, Alfredo Iacoangeli is funded by South London and Maudsley NHS Foundation Trust, MND Scotland, Motor Neuron Disease Association, National Institute for Health and Care Research, Spastic Paraplegia Foundation, Rosetrees Trust, Darby Rimmer MND Foundation, the Medical Research Council (UKRI), Alzheimer’s Research UK and LifeArc. The views expressed are those of the authors and not necessarily those of the NHS, the NIHR, the Department of Health and Social Care or the other funders.

## Data availability

Simulation trajectories are publicly deposited on Zenodo DOI: 10.5281/zenodo.20531241 and analysis scripts are available from the corresponding author upon reasonable request.

## 6. Supplementary

**Supplementary. S[1].**
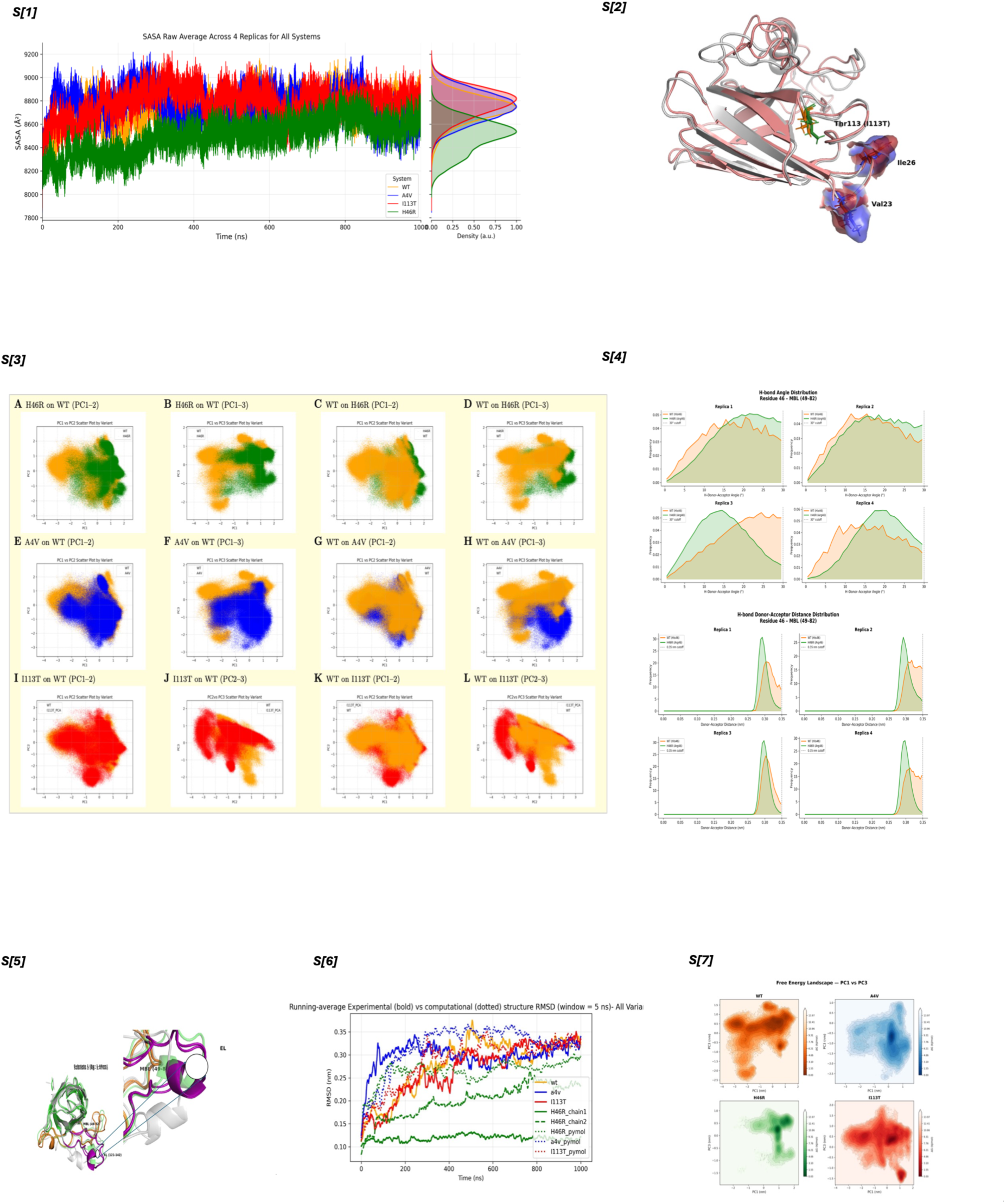
Solvent-accessible surface area (SASA)& Radius of gyration (Rg). All metrics are measured against the equilibrated structure averaged across n = 4 independent 1 μs replicas; right-hand panels show marginal kernel density estimates. Colour scheme: WT (orange), A4V (blue), H46R (green), I113T (red). H46R is the most compact and surface-minimised variant; I113T exhibits the greatest conformational heterogeneity across both measures. **S[2]**. **Structural comparison of WT and I113T SOD1 highlighting displacement of hydrophobic β-barrel residues Val23 and Ile26.** Superimposed representative structures of WT (grey cartoon) and I113T (salmon cartoon) SOD1 apo monomers aligned by backbone Cα atoms. Sticks and surface representations show residues Val23 and Ile26 (WT: blue; I113T: red) and the mutation site residue 113 (Ile113 WT: green; Thr113 I113T: orange). Insets show magnified views of the three residues of interest. The left inset illustrates the side-chain size reduction at position 113 upon Ile to Thr substitution, creating a cavity in the hydrophobic core. The right insets show the resulting displacement of Val23 and Ile26 side chains in I113T relative to WT, consistent with the elevated per-residue SASA observed at these positions (+26.2% and +38.6% respectively; Supplementary Figure S[1]). Together these structural observations support the hypothesis that the Ile113Thr substitution disrupts local hydrophobic packing, propagating conformational perturbation to neighbouring β-barrel residues and contributing to the elevated solvent exposure and conformational heterogeneity characteristic of I113T. **S[3].** Pairwise cross-projection of Cα MD trajectories onto shared PCA eigenvector bases. For each variant–WT pair, the trajectory of one system was projected onto the eigenvectors derived from the other (and vice versa), enabling direct comparison of conformational sampling within a shared coordinate frame. (A–D) H46R vs WT in the PC1–PC2 and PC1–PC3 planes. H46R samples conformational space shifted toward negative PC1 values relative to WT, with clearer separation in the PC1–PC3 projection, indicating altered electrostatic loop dynamics. (E–H) A4V vs WT in the PC1–PC2 and PC1–PC3 planes. Despite extensive overlap in PC1–PC2, the PC1–PC3 plane reveals that A4V accesses a distinct conformational basin at negative PC3 values not substantially populated by WT, consistent with novel loop rearrangements induced by the most clinically aggressive fALS mutation. (I–L) I113T vs WT in the PC1–PC2 and PC2–PC3 planes. I113T shows the greatest conformational overlap with WT among the three variants, with modest separation visible only in the PC2–PC3 projections. **S[4]**. H-bond geometry between residue 46 and MBL (residues 49–82). (A) Donor–acceptor distance distributions per replica; H46R (green) peaks at ∼0.29 nm, closer to the ideal H-bond distance than WT (orange, ∼0.30–0.31 nm). (B) H–donor–acceptor angle distributions; H46R peaks at lower angles (∼10–15°) than WT (∼15–20°), indicating more geometrically optimal H-bond geometry. Together with the contact data in Figure [1,2], these distributions confirm that Arg46 forms tighter and more directionally ideal interactions with the MBL than His46. **S[5]. H46R conformational substates identified from bimodal radius of gyration distribution figure S[1].** Superimposed representative structures extracted from the two Rg peaks of the H46R trajectory: substate 1 (Rg ∼1.46 nm; grey) and substate 2 (Rg ∼1.47 nm; green), with two independent replicas shown per substate (opaque and transparent respectively). The metal-binding loop (MBL, residues 49–82; orange) and electrostatic loop (EL, residues 121–142; purple) are highlighted. The right panel shows a magnified view of the loop region, revealing that the two substates differ predominantly in the position of the EL, while the MBL and β-barrel core remain largely invariant. This two-state EL behaviour is consistent with the bimodal EL RMSD distribution (Figure 2B) and the suppressed MBL–EL anti-correlated dynamics identified by DCCM (Figure 4C). **S[6]. Backbone RMSD of Experimental SOD1 structures vs Pymol induced point mutations. S[7]. FEL PC1 vs PC3 Projections S[7]. FEL PC1 vs PC3 Projections**

## References

(1) Pasinelli, P.; Brown, R. H. Molecular Biology of Amyotrophic Lateral Sclerosis: Insights from Genetics. Nat. Rev. Neurosci. 2006, 7 (9), 710–723. 10.1038/nrn1971.

(2) Masrori, P.; Van Damme, P. Amyotrophic Lateral Sclerosis: A Clinical Review. Eur. J. Neurol. 2020, 27 (10), 1918–1929. 10.1111/ene.14393.

(3) Andersen, P. M. Genetics of Sporadic ALS. Amyotroph. Lateral Scler. Other Motor Neuron Disord. 2001, 2 (SUPPL. 1), 37–41. 10.1080/14660820152415726.

(4) Rosen, D. R.; Siddique, T.; Patterson, D.; Figlewicz, D. A.; Sapp, P.; Hentati, A.; Donaldson, D.; Goto, J.; O’Regan, J. P.; Deng, H.-X.; Rahmani, Z.; Krizus, A.; McKenna-Yasek, D.; Cayabyab, A.; Gaston, S. M.; Berger, R.; Tanzi, R. E.; Halperin, J. J.; Herzfeldt, B.; Den Bergh, R. V.; Hung, W.-Y.; Bird, T.; Deng, G.; Mulder, D. W.; Smyth, C.; Laing, N. G.; Soriano, E.; Pericak-Vance, M. A.; Haines, J.; Rouleau, G. A.; Gusella, J. S.; Horvitz, H. R.; Brown Jr., R. H. Mutations in Cu/Zn Superoxide Dismutase Gene Are Associated with Familial Amyotrophic Lateral Sclerosis. Nature 1993, 362 (6415), 59–62. 10.1038/362059a0.

(5) Ghasemi, R. H., Mehdi ;. Brown. Genetics of Amyotrophic Lateral Sclerosis. Cold Spring Harb. Perspect. Med. 2018, 8. 10.1101/cshperspect.a024125.

(6) Valentine, J. S.; Doucette, P. A.; Potter, S. Z. COPPER-ZINC SUPEROXIDE DISMUTASE AND AMYOTROPHIC LATERAL SCLEROSIS. Annu. Rev. Biochem. 2005, 74 (Volume 74, 2005), 563–593. 10.1146/annurev.biochem.72.121801.161647.

(7) Arnesano, F.; Banci, L.; Bertini, I.; Martinelli, M.; Furukawa, Y.; O’Halloran, T. V. The Unusually Stable Quaternary Structure of Human Cu,Zn-Superoxide Dismutase 1 Is Controlled by Both Metal Occupancy and Disulfide Status*. J. Biol. Chem. 2004, 279 (46), 47998–48003. 10.1074/jbc.M406021200.

(8) Getzoff, E. D.; Tainer, J. A.; Stempien, M. M.; Bell, G. I.; Hallewell, R. A. Evolution of CuZn Superoxide Dismutase and the Greek Key β-Barrel Structural Motif. Proteins Struct. Funct. Bioinforma. 1989, 5 (4), 322–336. 10.1002/prot.340050408.

(9) Galaleldeen, A.; Strange, R. W.; Whitson, L. J.; Antonyuk, S. V.; Narayana, N.; Taylor, A. B.; Schuermann, J. P.; Holloway, S. P.; Hasnain, S. S.; Hart, P. J. Structural and Biophysical Properties of Metal-Free Pathogenic SOD1 Mutants A4V and G93A. Arch. Biochem. Biophys. 2009, 492 (1), 40–47. 10.1016/j.abb.2009.09.020.

(10) Nedd, S.; Redler, R. L.; Proctor, E. A.; Dokholyan, N. V.; Alexandrova, A. N. Cu, Zn-Superoxide Dismutase without Zn Is Folded but Catalytically Inactive. J. Mol. Biol. 2014, 426 (24), 4112–4124. 10.1016/j.jmb.2014.07.016.

(11) Furukawa, Y.; Anzai, I.; Akiyama, S.; Imai, M.; Cruz, F. J. C.; Saio, T.; Nagasawa, K.; Nomura, T.; Ishimori, K. Conformational Disorder of the Most Immature Cu, Zn-Superoxide Dismutase Leading to Amyotrophic Lateral Sclerosis. J. Biol. Chem. 2016, 291 (8), 4144–4155. 10.1074/jbc.M115.683763.

(12) Furukawa, Y. Good and Bad of Cu/Zn-Superoxide Dismutase Controlled by Metal Ions and Disulfide Bonds. Chem. Lett. 2021, 50 (2), 331–341. 10.1246/cl.200770.

(13) Andersen, P. M.; Al-Chalabi, A. Clinical Genetics of Amyotrophic Lateral Sclerosis: What Do We Really Know? Nat. Rev. Neurol. 2011, 7 (11), 603–615. 10.1038/nrneurol.2011.150.

(14) Berdyñski, M.; Miszta, P.; Safranow, K.; Andersen, P. M.; Morita, M.; Filipek, S.; Zekanowski, C.; Kuźma-Kozakiewicz, M. SOD1 Mutations Associated with Amyotrophic Lateral Sclerosis Analysis of Variant Severity. Sci. Rep. 2022, 12 (1). 10.1038/s41598-021-03891-8.

(15) Kalia, M.; Miotto, M.; Ness, D.; Opie-Martin, S.; Spargo, T. P.; Di Rienzo, L.; Biagini, T.; Petrizzelli, F.; Al Khleifat, A.; Kabiljo, R.; Project MinE ALS Sequencing Consortium; SOD1-ALS clinical and genetic data collection group; Mazza, T.; Ruocco, G.; Milanetti, E.; Dobson, R. J.; Al-Chalabi, A.; Iacoangeli, A. Molecular Dynamics Analysis of Superoxide Dismutase 1 Mutations Suggests Decoupling between Mechanisms Underlying ALS Onset and Progression. Comput. Struct. Biotechnol. J. 2023, 21, 5296–5308. 10.1016/j.csbj.2023.09.016.

(16) Basith, S.; Manavalan, B.; Lee, G. Amyotrophic Lateral Sclerosis Disease-Related Mutations Disrupt the Dimerization of Superoxide Dismutase 1 - A Comparative Molecular Dynamics Simulation Study. Comput. Biol. Med. 2022, 151. 10.1016/j.compbiomed.2022.106319.

(17) Jahan, I.; Nayeem, S. M. Conformational Dynamics of Superoxide Dismutase (SOD1) in Osmolytes: A Molecular Dynamics Simulation Study. RSC Adv. 2020, 10 (46), 27598–27614. 10.1039/D0RA02151B.

(18) Vassall, K. A.; Stathopulos, P. B.; Rumfeldt, J. A. O.; Lepock, J. R.; Meiering, E. M. Equilibrium Thermodynamic Analysis of Amyotrophic Lateral Sclerosis-Associated Mutant Apo Cu,Zn Superoxide Dismutases,. Biochemistry 2006, 45 (23), 7366–7379. 10.1021/bi0600953.

(19) Keskin, I.; Forsgren, E.; Lehmann, M.; Andersen, P. M.; Brännström, T.; Lange, D. J.; Synofzik, M.; Nordström, U.; Zetterström, P.; Marklund, S. L.; Gilthorpe, J. D. The Molecular Pathogenesis of Superoxide Dismutase 1-Linked ALS Is Promoted by Low Oxygen Tension. Acta Neuropathol. (Berl.) 2019, 138 (1), 85–101. 10.1007/s00401-019-01986-1.

(20) Strange, R. W.; Antonyuk, S. V.; Hough, M. A.; Doucette, P. A.; Valentine, J. S.; Hasnain, S. S. Variable Metallation of Human Superoxide Dismutase: Atomic Resolution Crystal Structures of Cu–Zn, Zn–Zn and As-Isolated Wild-Type Enzymes. J. Mol. Biol. 2006, 356 (5), 1152–1162. 10.1016/j.jmb.2005.11.081.

(21) Hough, M. A.; Grossmann, J. G.; Antonyuk, S. V.; Strange, R. W.; Doucette, P. A.; Rodriguez, J. A.; Whitson, L. J.; Hart, P. J.; Hayward, L. J.; Valentine, J. S.; Hasnain, S. S. Dimer Destabilization in Superoxide Dismutase May Result in Disease-Causing Properties: Structures of Motor Neuron Disease Mutants. Proc. Natl. Acad. Sci. 2004, 101 (16), 5976–5981. 10.1073/pnas.0305143101.

(22) Elam, J. S.; Taylor, A. B.; Strange, R.; Antonyuk, S.; Doucette, P. A.; Rodriguez, J. A.; Hasnain, S. S.; Hayward, L. J.; Valentine, J. S.; Yeates, T. O.; Hart, P. J. Amyloid-like Filaments and Water-Filled Nanotubes Formed by SOD1 Mutant Proteins Linked to Familial ALS. Nat. Struct. Mol. Biol. 2003, 10 (6), 461–467. 10.1038/nsb935.

(23) Hough, M. A.; Grossmann, J. G.; Antonyuk, S. V.; Strange, R. W.; Doucette, P. A.; Rodriguez, J. A.; Whitson, L. J.; Hart, P. J.; Hayward, L. J.; Valentine, J. S.; Hasnain, S. S. I113T Mutant of Human SOD1: 1uxl, 2004. 10.2210/pdb1uxl/pdb.

(24) PyMOL Molecular Graphics System - Browse /pymol/1.8 at SourceForge.net. https://sourceforge.net/projects/pymol/files/pymol/1.8/ (accessed 2026-05-19).

(25) Van Der Spoel, D.; Lindahl, E.; Hess, B.; Groenhof, G.; Mark, A. E.; Berendsen, H. J. C. GROMACS: Fast, Flexible, and Free. J. Comput. Chem. 2005, 26 (16), 1701–1718. 10.1002/jcc.20291.

(26) Lindahl; Abraham; Hess; Spoel, van der. GROMACS 2021.5 Source Code, 2022. 10.5281/zenodo.5850051.

(27) Mackerell, A. D.; Feig, M.; Brooks, C. L. Extending the Treatment of Backbone Energetics in Protein Force Fields: Limitations of Gas-Phase Quantum Mechanics in Reproducing Protein Conformational Distributions in Molecular Dynamics Simulations. J. Comput. Chem. 2004, 25 (11), 1400–1415. 10.1002/jcc.20065.

(28) MacKerell, A. D. Jr.; Bashford, D.; Bellott, M.; Dunbrack, R. L. Jr.; Evanseck, J. D.; Field, M. J.; Fischer, S.; Gao, J.; Guo, H.; Ha, S.; Joseph-McCarthy, D.; Kuchnir, L.; Kuczera, K.; Lau, F. T. K.; Mattos, C.; Michnick, S.; Ngo, T.; Nguyen, D. T.; Prodhom, B.; Reiher, W. E.; Roux, B.; Schlenkrich, M.; Smith, J. C.; Stote, R.; Straub, J.; Watanabe, M.; Wiórkiewicz-Kuczera, J.; Yin, D.; Karplus, M. All-Atom Empirical Potential for Molecular Modeling and Dynamics Studies of Proteins. J. Phys. Chem. B 1998, 102 (18), 3586–3616. 10.1021/jp973084f.

(29) Freddolino, L.; Liu, F.; Gruebele, M.; Schulten, K. Ten-Microsecond Molecular Dynamics Simulation of a Fast-Folding WW Domain. Biophys. J. 2008, 94 (10), L75–L77. 10.1529/biophysj.108.131565.

(30) Berendsen, H. J. C.; Grigera, J. R.; Straatsma, T. P. The Missing Term in Effective Pair Potentials. J. Phys. Chem. 1987, 91 (24), 6269–6271. 10.1021/j100308a038.

(31) Essmann, U.; Perera, L.; Berkowitz, M. L.; Darden, T.; Lee, H.; Pedersen, L. G. A Smooth Particle Mesh Ewald Method. J. Chem. Phys. 1995, 103 (19), 8577–8593. 10.1063/1.470117.

(32) Berendsen, H. J. C.; Postma, J. P. M.; van Gunsteren, W. F.; DiNola, A.; Haak, J. R. Molecular Dynamics with Coupling to an External Bath. J. Chem. Phys. 1984, 81 (8), 3684–3690. 10.1063/1.448118.

(33) Parrinello, M.; Rahman, A. Polymorphic Transitions in Single Crystals: A New Molecular Dynamics Method. J. Appl. Phys. 1981, 52 (12), 7182–7190. 10.1063/1.328693.

(34) Hess, B.; Bekker, H.; Berendsen, H. J. C.; Fraaije, J. G. E. M. LINCS: A Linear Constraint Solver for Molecular Simulations. J. Comput. Chem. 1997, 18 (12), 1463–1472. 10.1002/(SICI)1096-987X(199709)18:12%3C1463::AID-JCC4%3E3.0.CO;2-H.

(35) Matplotlib: A 2D Graphics Environment | IEEE Journals & Magazine | IEEE Xplore. https://ieeexplore.ieee.org/document/4160265 (accessed 2026-05-20).

(36) Grant, B. J.; Rodrigues, A. P. C.; ElSawy, K. M.; McCammon, J. A.; Caves, L. S. D. Bio3d: An R Package for the Comparative Analysis of Protein Structures. Bioinformatics 2006, 22 (21), 2695–2696. 10.1093/bioinformatics/btl461.

(37) McKinney, W. Data Structures for Statistical Computing in Python. SciPy 2010 2010. 10.25080/Majora-92bf1922-00a.

(38) Harris, C. R.; Millman, K. J.; van der Walt, S. J.; Gommers, R.; Virtanen, P.; Cournapeau, D.; Wieser, E.; Taylor, J.; Berg, S.; Smith, N. J.; Kern, R.; Picus, M.; Hoyer, S.; van Kerkwijk, M. H.; Brett, M.; Haldane, A.; del Río, J. F.; Wiebe, M.; Peterson, P.; Gérard-Marchant, P.; Sheppard, K.; Reddy, T.; Weckesser, W.; Abbasi, H.; Gohlke, C.; Oliphant, T. E. Array Programming with NumPy. Nature 2020, 585 (7825), 357–362. 10.1038/s41586-020-2649-2.

(39) Huai, J.; Zhang, Z. Structural Properties and Interaction Partners of Familial ALS-Associated SOD1 Mutants. Front. Neurol. 2019, 10. 10.3389/fneur.2019.00527.

(40) Kim, J.; Lee, H.; Lee, J. H.; Kwon, D.-Y.; Genovesio, A.; Fenistein, D.; Ogier, A.; Brondani, V.; Grailhe, R. Dimerization, Oligomerization, and Aggregation of Human Amyotrophic Lateral Sclerosis Copper/Zinc Superoxide Dismutase 1 Protein Mutant Forms in Live Cells. J. Biol. Chem. 2014, 289 (21), 15094–15103. 10.1074/jbc.M113.542613.

(41) Perri, E. R.; Parakh, S.; Vidal, M.; Mehta, P.; Ma, Y.; Walker, A. K.; Atkin, J. D. The Cysteine (Cys) Residues Cys-6 and Cys-111 in Mutant Superoxide Dismutase 1 (SOD1) A4V Are Required for Induction of Endoplasmic Reticulum Stress in Amyotrophic Lateral Sclerosis. J. Mol. Neurosci. 2020, 70 (9), 1357–1368. 10.1007/s12031-020-01551-6.

(42) Basith, S.; Manavalan, B.; Lee, G. Unveiling Local and Global Conformational Changes and Allosteric Communications in SOD1 Systems Using Molecular Dynamics Simulation and Network Analyses. Comput. Biol. Med. 2024, 168. 10.1016/j.compbiomed.2023.107688.

(43) Nordlund, A.; Oliveberg, M. SOD1-Associated ALS: A Promising System for Elucidating the Origin of Protein-Misfolding Disease. HFSP J. 2008, 2 (6), 354–364. 10.2976/1.2995726.

(44) Rasouli, S.; Abdolvahabi, A.; Croom, C. M.; Plewman, D. L.; Shi, Y.; Shaw, B. F. Glycerolipid Headgroups Control Rate and Mechanism of Superoxide Dismutase-1 Aggregation and Accelerate Fibrillization of Slowly Aggregating Amyotrophic Lateral Sclerosis Mutants. ACS Chem. Neurosci. 2018, 9 (7), 1743–1756.

(45) Eastman, J. Swails, J. D. Chodera, R. T. McGibbon, Y. Zhao, K. A. Beauchamp, L.-P. Wang, A. C. Simmonett, M. P. Harrigan, C. D. Stern, R. P. Wiewiora, B. R. Brooks, an d V. S. Pande. “OpenMM 7: Rapid development of high performance algorithms for molecular dynamics.” PLOS Comp. Biol. 13(7): e1005659. (2017)

